# Effects of Cholesterol on the mechanism of fengycin, a biofungicide

**DOI:** 10.1101/2021.08.02.454812

**Authors:** Sreyoshi Sur, Alan Grossfield

## Abstract

Fengycins are a class of antifungal lipopeptides synthesized by the bacteria *Bacillus subtilis*, commercially available as the primary component of the agricultural fungicide Serenade^®^. They are toxic to fungi, but far less to mammalian cells. One key difference between mammalian and fungal cell membranes is the presence of cholesterol only in the former; recent experimental work showed that the presence of cholesterol reduces fengycin-induced membrane leakage. (1) Since our previous all-atom and coarse-grained simulations (2, 3) suggested that aggregation of membrane-bound fengycin is central to its ability to disrupt membranes, we hypothesized that cholesterol might reduce fengycin aggregation. Here, we test this hypothesis using coarse-grained molecular dynamics simulations, with sampling enhanced via the weighted ensemble method. The results indicate that cholesterol subtly alters the size distribution for fengycin aggregates, limits the lateral range of their membrane disordering, and reduces the ability of aggregates to bend the membrane. Taken together, these phenomena may account for cholesterol’s affects on fengycin activity.

## 1 Introduction

The world human population has reached 7.8 billion as of August 2021 (4), which creates ever greater pressure to increase agricultural yields.(5) Every year crops are destroyed by various microbial infections, of which 60% are caused by fungus.(6) To cope with this, various antibiotics have been synthesized like amphotericin B, natamycin and nystatin(7–9) but fungi and bacteria are developing resistance to these compounds.

With this in mind, many groups are interested in developing new ecologically friendly antifungals suitable for agricultural usage.(10) Fengycins are a class of lipopeptides synthesized by the bacteria *Bacillus subtilis*, which specifically kill certain fungal species.(11, 12) Fengycins, along with surfactins and iturins, are the primary active components of Serenade^®^ (Bayer), a commercial fungicide and bacteriocide, which is used to treat plant diseases like clubroot disease (*Plasmodiophora monoliforme*), barley head blight (*Fusarium graminearum*), cucurbit powdery (*Podosphaera fusca*), black Sigatoka (*Mycosphaerella fijiensis*), grey mold (*Botrytis cinerea*) and soft rot (*Rhizopus stolonifer*).(13–22)

The effects of fengycin seem specific to fungal cells, with far less effect on animals. This is part of its “green” signature, since neither the agricultural workers applying it nor the animals in the surrounding environment are adversely affected by its application. It would be advantageous to understand why this is the case, particularly given that it has drawn medical interest as a potential anti-fungal skin ointment. (23–25)

Previous calorimetric studies suggested that fengycins cause order to disorder phase transitions and partition into the cholesterol rich phase in the stratum corneum (outermost skin layer) model and in monolayers mimicking them.(26, 27) This implies that either fengycin prefers a more lipid disordered phase or that it preferentially interacts with cholesterol.

To better understand the mechanism of fengycin’s selectivity, Heerklotz and coworkers conducted fluorescence lifetime experiments with model membranes of varying phospholipids and sterols composition.(28, 29) They showed that higher concentrations of fengycin were required to induce leakage when the vesicles contain cholesterol, indicating that cholesterol inhibits fengycin. The authors discussed two possible mechanisms for this phenomenon: a) graded leakage, where the fengycin’s induce general overall leakiness by thinning of membrane and disordering the lipids, or b) all-or-none leakage, where long-lived pores form which cause the vesicles to drain quickly and completely.(30) Patel et al. showed that fengycin follows an all-or-none mechanism using these methods, (30) while Fiedler et al. found that the tolerable fraction of fengycins on the membrane surface increases in a mixture of POPC and cholesterol membranes compared to pure POPC.(1) The purported presence of pores suggests that fengycin must aggregate on the membrane surface in order to function, and although no experiments have been able to demonstrate this effect directly, Fiedler et al hypothesized that fengycins preferentially interact with cholesterols and thus are unable to aggregate, reducing membrane permeabilization.(31)

Molecular dynamics simulations are an excellent tool to fill in this missing connection, since their molecule-scale resolution neatly matches the problem. However, obtaining sufficient sampling still remains an issue; lateral reorganization of the membrane and protein conformational change are both slow processes compared to the timescale of most molecular dynamics simulations. Use of coarse-grained simulations reduces the computational cost of sampling, as does our application of the weighted ensemble method.(32, 33)

Previous work from our group showed that fengycin aggregation was higher in a zwitterionic lipid membrane chosen to model eukaryotes, compared to anionic membranes chosen to mimic bacteria, and that aggregation was crucial to disordering the membrane.(2, 3) Here, we study a more subtle variation, the effect of cholesterol on fengycin aggregation in the backdrop of simple zwitterionic bilayer.

## 2 Materials and Methods

### 2.1 System construction

All the particles in our systems were modeled using the MARTINI coarse-grained force field ver. 2.2P.(34) We considered two lipid compositions: pure palmitoyl-oleoyl-phosphatidylcholine (POPC), and a 4:1 mixture of POPC and cholesterol. The POPC systems contained 186 POPC molecules, with 8352 waters, and 99 Na+ and 84 Clions. Each composition was run with 15 fengycin molecules per leaflet; neat lipid simulations were run for comparison purposes. The systems containing fengycin contained an additional 15 sodium ions to maintain electroneutrality. The parameters for fengycin were adapted from Horn et al. (2), after accounting for force field updates. For lipids, we used MARTINI Ver. 2.0, and for the peptide part of fengycin MARTINI Ver. 2.2 was used.(34) The parameters are available in our github repository. The chemical structure of fengycin is shown in Figure 1A; this variant is also referred to as Plipastatin A in the literature. The sequence differs slightly from the one we simulated previously, with glutamate 8 replaced with glutamine. (2)

**Figure 1:**
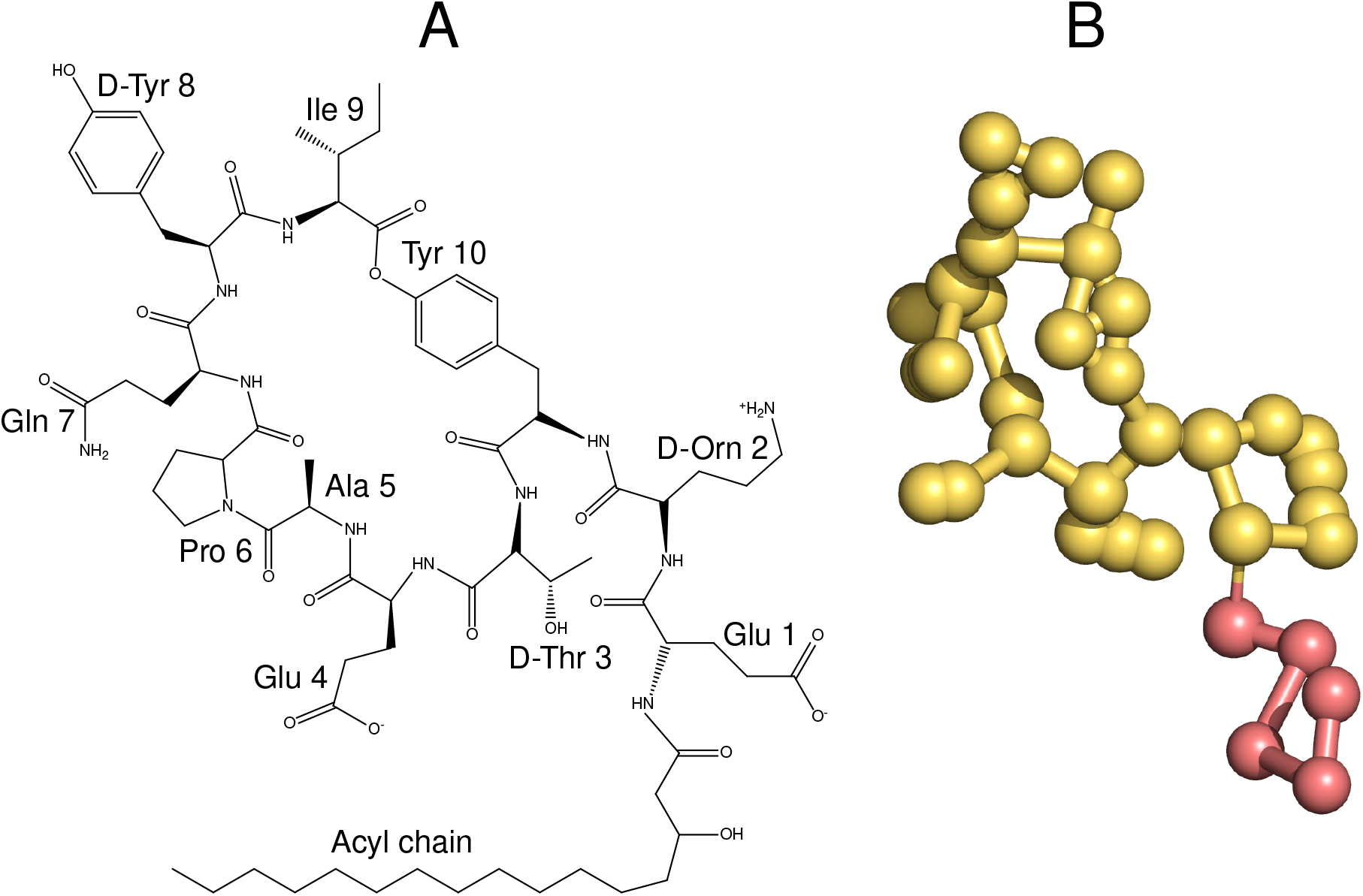
A. Chemical structure of fengycin B.Coarse-grained model of fengycin.

Initially, we randomly placed fengycins in a plane parallel to the bilayer, with each fengycins 20 Å above the bilayer center and the acyl chain pointing toward the membrane center. We constructed the rest of the membrane around them using INSANE,(35) and added NaCl for an effective salt concentration of 150 mM using GROMACS genion tool.(36) Next, we relaxed each system with 10000 steps of energy minimization, followed by 10-15 ns of molecular dynamics simulations with a timestep of 2 ns. All calculations were performed using GROMACS Ver. 2016.3.(36) The Table 1 shows all the details about the systems we built.

**Table 1:** Summary of simulations. “WE” refers to weighted ensemble, while “MD” indicates standard molecular dynamics. For the WE runs, the time refers to the total sampling time for each WE replicate. “Replicates” the number of independently built trajectories that were run; 3 distinct WE simulations were performed for each composition.

| Membrane | Peptides | Length(μs) | Replicates | Method |
| --- | --- | --- | --- | --- |
| PC | 15 | ∼350 | 3 | WE |
| PC:CHOL | 15 | ∼ 330 | 3 | WE |
| PC | 0 | ∼0.6 | 3 | MD |
| PC:CHOL | 0 | ∼0.6 | 3 | MD |

### 2.2 Simulation details

We used the Gromacs Ver. 2016.3 to perform the molecular dynamics simulations.(36) We used a timestep of 20 fs, updating the nearest neighbor list every 20 steps. The simulations were run in an isobaric-isothermic ensemble with the Parrinello-Rahman barostat (37, 38) and the velocity rescaling thermostat.(39) Electrostatic interactions were calculated using the reaction field method.(40) The Verlet scheme was used to compute the Van der waals forces using a 11 Å cutoff. The cumulative time of the simulation is shown in Table 1.

### 2.3 Weighted ensemble (WE) method

We used the weighted ensemble method to calculate the probability distribution for fengycin aggregation,(41) implemented via the WESTPA software package (42, 43). In this method, no additional bias or force is added to the system to enhance the sampling. Rather, structures are binned along a pre-chosen progress coordinate, and trajectories are split and merged to maintain a target number of trajectories in each populated bin, with the overall statistical weight tracked appropriately.

We quantify the aggregation of fengycin by using as progress coordinate the total number of contacts between fengycin molecules. The contact value for a pair of fengycins i and j is:

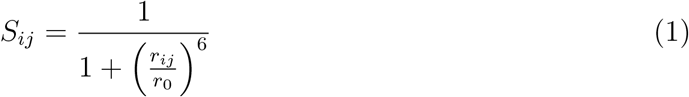

where *r*_*ij*_ is the lateral distance between the centroids of the peptide part of fengycins and *r*_0_ = 10Å. The total number of contacts for a given frame in the trajectory is the sum of *S*_*ij*_ between all pairs of fengycins,

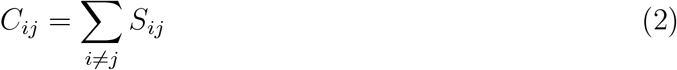

This progress coordinate was divided into 25 evenly spaced bins each bin having a width of 1.0.

Each iteration of weighted ensemble simulations involves two stages. First, each starting structure is subjected to 5 ns of molecular dynamics simulations. At the end of the dynamics, each trajectory is assigned to a bin based on its total contact value (*C*_*ij*_, from Equation 2). We chose to target 5 trajectories to each populated bin. If there were more than 5 in a given bin, pairwise approach is used to prune a trajectory. Within the 5 trajectories two are chosen at random of which only one with relatively less weight is pruned and the weight of the pruned trajectory is assigned to the surviving trajectory. If there were fewer than 5 trajectories, a trajectory was chosen at random and split, with its weight equally shared with the resulting children. The total weight of all the trajectories for a particular iteration should always be 1.0.(33, 42)

### 2.4 Simulation analysis

LOOS was used to analyze all the data, along with the analysis tools that came with the WESTPA software (42, 44–46). All the analysis was performed at 1 ns resolution. All average quantities were computed using the last 50 WE iterations, taking into account the weight associated with each trajectory. Unless otherwise specified, error bars are the standard errors calculated by treating each weighted ensemble simulation as one independent measurement.

#### 2.4.1 Molecular order parameters

The molecular order parameter is a whole-chain analog of the deuterium order parameters measured using solid-state NMR.(47) It is calculated as

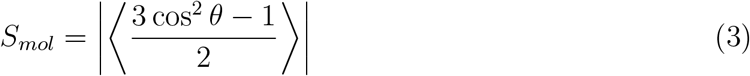

where *θ* is the angle between the membrane normal and the second and third principal axes of the acyl chain; this method is implemented in the *dibmops* tool in LOOS.

#### 2.4.2 Radial Distribution function (RDF)

We used the LOOS tool *xy_rdf* to calculate the two-dimensional lateral radial distribution function in the plane of the membrane for various membrane component (lipid, cholesterol, fengycin); this tool operates on the molecules’ centroid, rather than any individual atoms.

#### 2.4.3 Probability distribution for fengycin-fengycin contacts and fengycin-lipid contacts

We used the tool *w_pdist* from WESTPA to compare weighted population distributions along the progress coordinate for each WE iteration. *plothist* (also part of WESTPA) was used to plot time series of this distribution, as well as averages over specific ranges of iterations.

#### 2.4.4 Aggregation propensity

When quantifying aggregation, we considered two fengycins to be contact if they have 50 pairs of atoms less than 6.5 Å apart; these specific values were chosen to reproduce manual assignments made by eye in a subset of trial frames. Using this criterion, we tracked the size distribution of fengycin aggregates through the weighted ensemble runs. As above, we used *w_pdist* and *plothist* to create weighted histograms for the number fengycin-fengycin contacts and the total number of aggregates. (42, 43)

#### 2.4.5 Z-position of phosphate heads

We tracked the deformation of the membrane by calculating the z-position of the phosphate bead of each POPC with respect to the center of membrane. These distances were binned with respect to the lateral distance to the nearest fengycin’s centroid. To simplify comparison between systems with and without cholesterol, we subtracted the thickness of the corresponding pure membranes to measure the perturbation caused due to fengycin.

All the analysis scripts and the MARTINI parameter files are available as a github repository at https://github.com/sreyoshi09/scripts.git

## 3 Results

Our previous work showed that larger aggregates, survive for a longer lifetime in POPC bilayers compared to POPE:POPG ones.(3)

Previous experimental work indicated that cholesterol reduces the ability of fengycin to induce membrane leakage, and hypothesized the presence of specific cholesterol-fengycin interactions to account for it.(31) Here, we test an alternative hypothesis, that cholesterol changes the aggregation of fengycin, such that it is less able to induce membrane deformation and leakage.

### 3.1 Fengycin forms ordered aggregates

Figure 2A shows the lateral radial distribution function for fengycin molecules. Large first peak indicates a strong tendency to oligomerize, while numerous secondary peaks indicate the presence of higher order structures. It seems that fengycin has a tendency to form some solid-gel systems as evident from their solid-like radial distribution function. Furthermore, the fact that the peaks are equally spaced, well-resolved, and of similar width strongly suggests that the aggregates are linear, a fact we verified visually (see Figure 2B). Although previous coarse-grained simulations also showed the presence of linear aggregates, our more recent all-atom simulations suggest the aggregates were more amorphous, consistent with atomic force microscopy experiments from Eeman et al.(2, 3, 25) It is worth noting that the MARTINI coarse-grained model is known to overestimate the aggregation propensity of proteins in MARTINI model (48).

**Figure 2:**
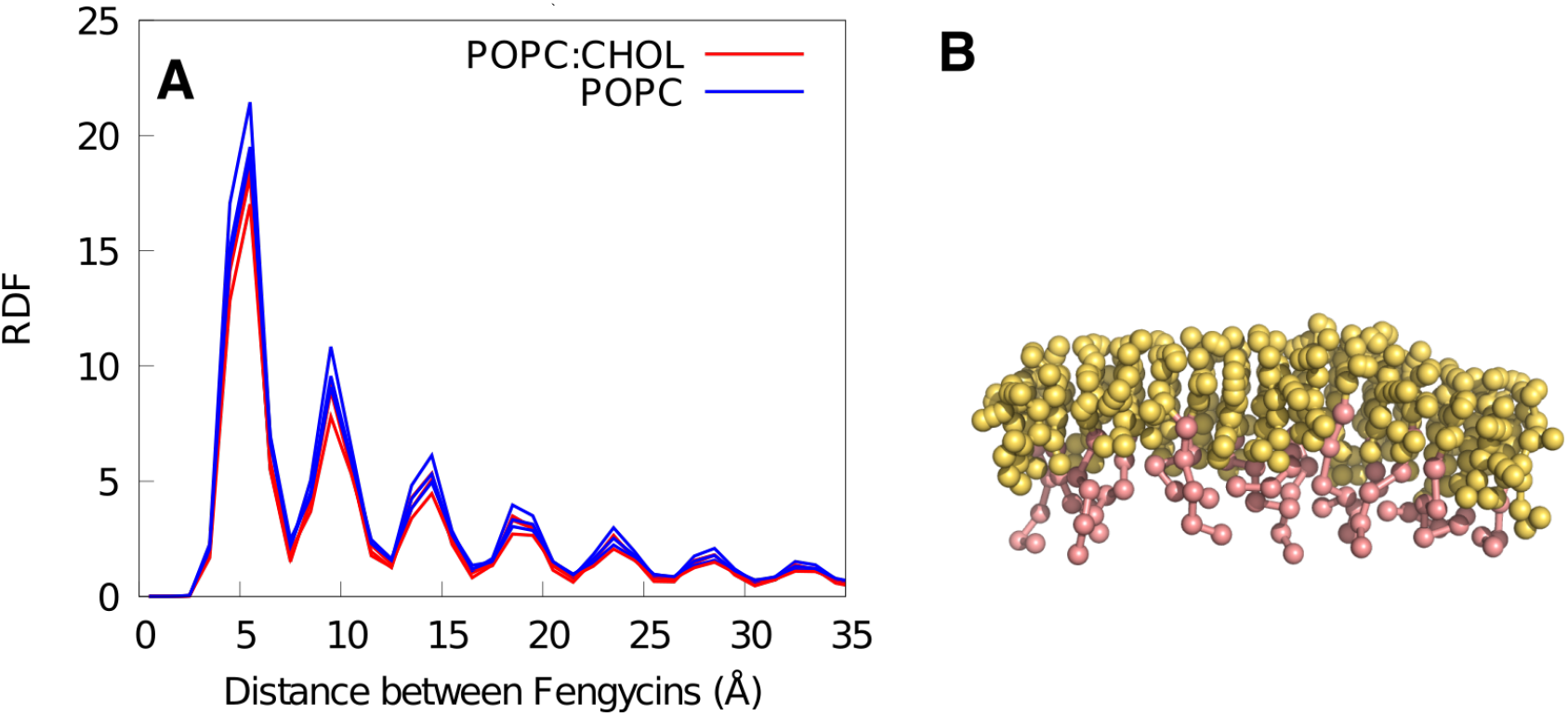
A. Lateral radial distribution function between fengycins. Each curve represents the average from a single weighted ensemble run. B. Structure of a single 15-mer aggregate

### 3.2 Aggregation propensity higher in cholesterol

Fiedler et al.’s fluorescence lifetime experiments suggested that cholesterol reduces fengycin-induced membrane leakage by reducing fengycin’s ability to aggregate in the membrane(31). One of the primary goals of the present work is to test this hypothesis.

As discussed in Section 2.4.3, we track aggregation by counting the number of fengycin-fengycin contacts; the resulting probability distribution is shown as Figure 3. Somewhat surprisingly, the data indicates that the presence of cholesterol slightly increases the tendency of fengycin to aggregate. Although the peak of the probability distribution is virtually identical, the tail to lower numbers of contacts is wider in the membrane without cholesterol, which also contains a measurable population of fully dispersed fengycin molecules.

**Figure 3:**
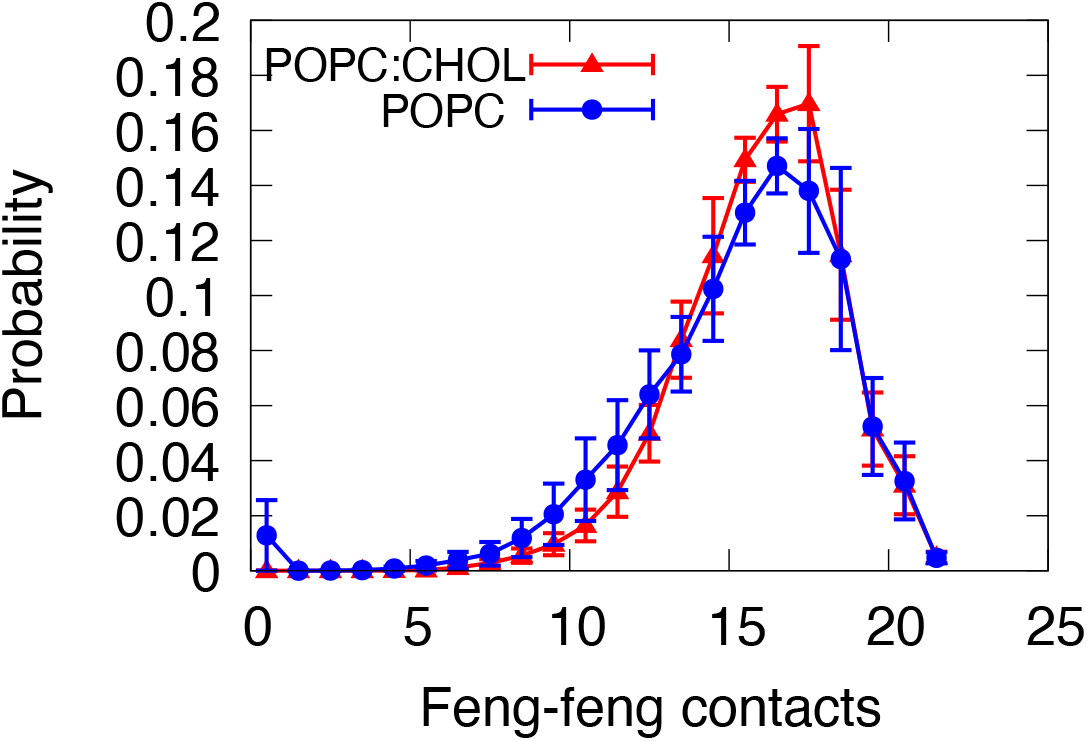
Probability distribution for fengycin-fengycin contacts. The number of fengycin contacts is calculated using Equations 1 and 2. Error bars are the standard error, calculated by considering each weighted ensemble run as a single independent measurement.

Reducing the complex phenomenon of aggregation to a single progress coordinate necessarily reduces the amount of information available. To recover some of this information, we also track a 2-dimensional probability distribution, as a function of number of contacts and the number of distinct aggregates, which allows us to distinguish potential changes in the morphology of the clusters.

The results are shown in Figure 4. Panels A and B show the distributions in the POPC and POPC:cholesterol membranes, respectively. Since the two distributions are fairly similar, we show the difference between them as Panel C. Focusing on Panel C, it is clear that the fengycins are more likely to be found as a single large aggregate in the presence of cholesterol, while in a pure POPC membrane there is more diversity in both the number of contacts and number of distinct aggregates.

**Figure 4:**
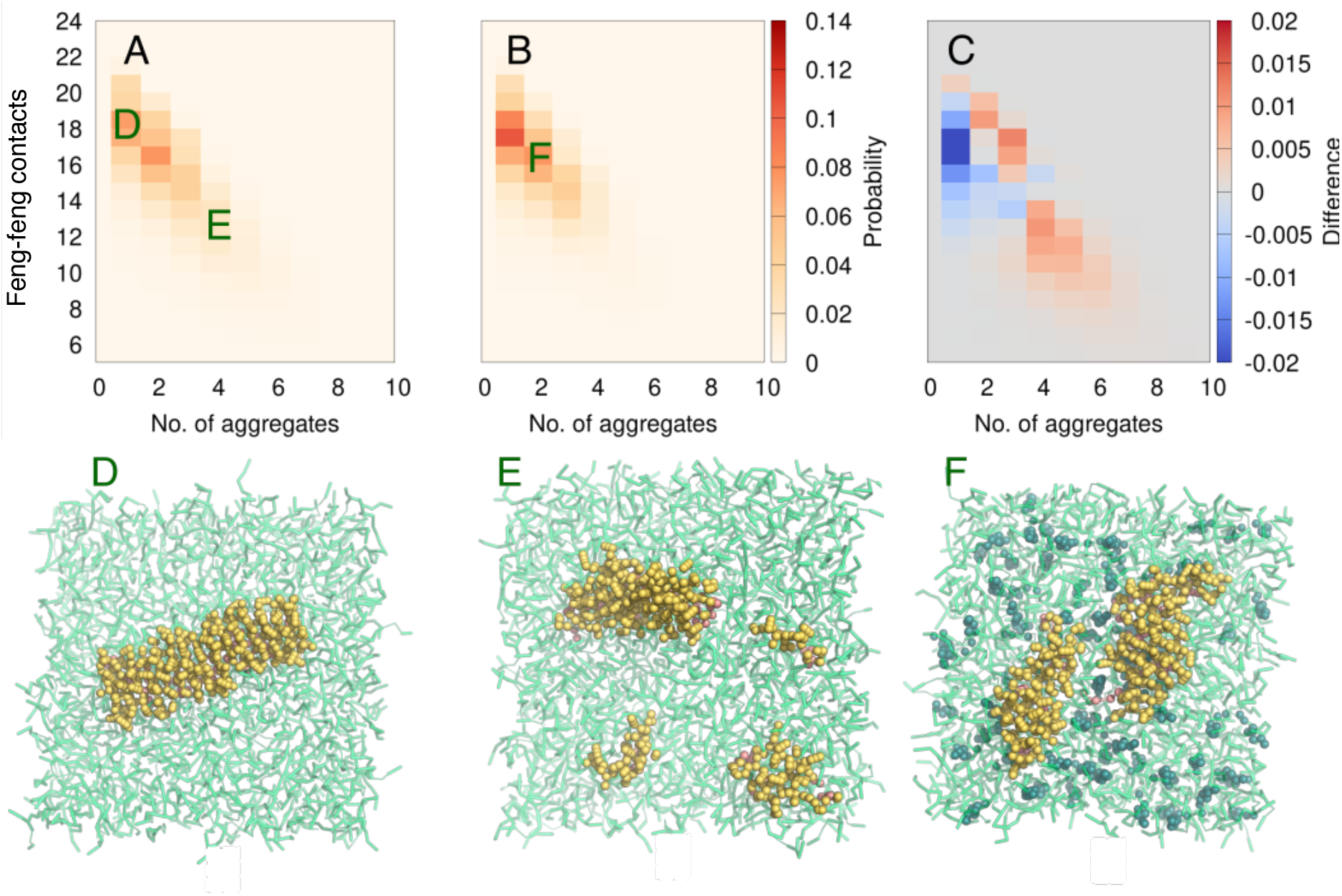
Probability of number of feng-feng contacts and number of aggregates in A. POPC and B. POPC:CHOL. C is obtained by subtracting the probabilities of B from A (POPC - POPC:CHOL). Example structures from different part of the probability distribution are shown in Panels D, E, and F; their locations on the 2D projection are indicated as green letters superposed onto Panels A and B.

Panels D, E, and F of Figure 4 indicate the kinds of aggregation consistent with specific parts of the probability distribution. Most notably, Panel E shows that when we have relatively high numbers of contacts *and* numbers of aggregates, this generally indicates the presence of one or two major aggregates, plus a number of isolated monomers.

### 3.3 Fengycin does not specifically pack with cholesterol or phospholipids

Our previous all-atom simulations suggested that fengycin aggregation was in part due to lack of favorable interactions with lipids, as opposed to strong specific attraction between the fengycins themselves.

Here, we extend that analysis to check for preferential localization with POPC and cholesterol. Specifically, we plotted the lateral radial distribution function between fengycin and cholesterol or POPC, shown in Figure5.

**Figure 5:**
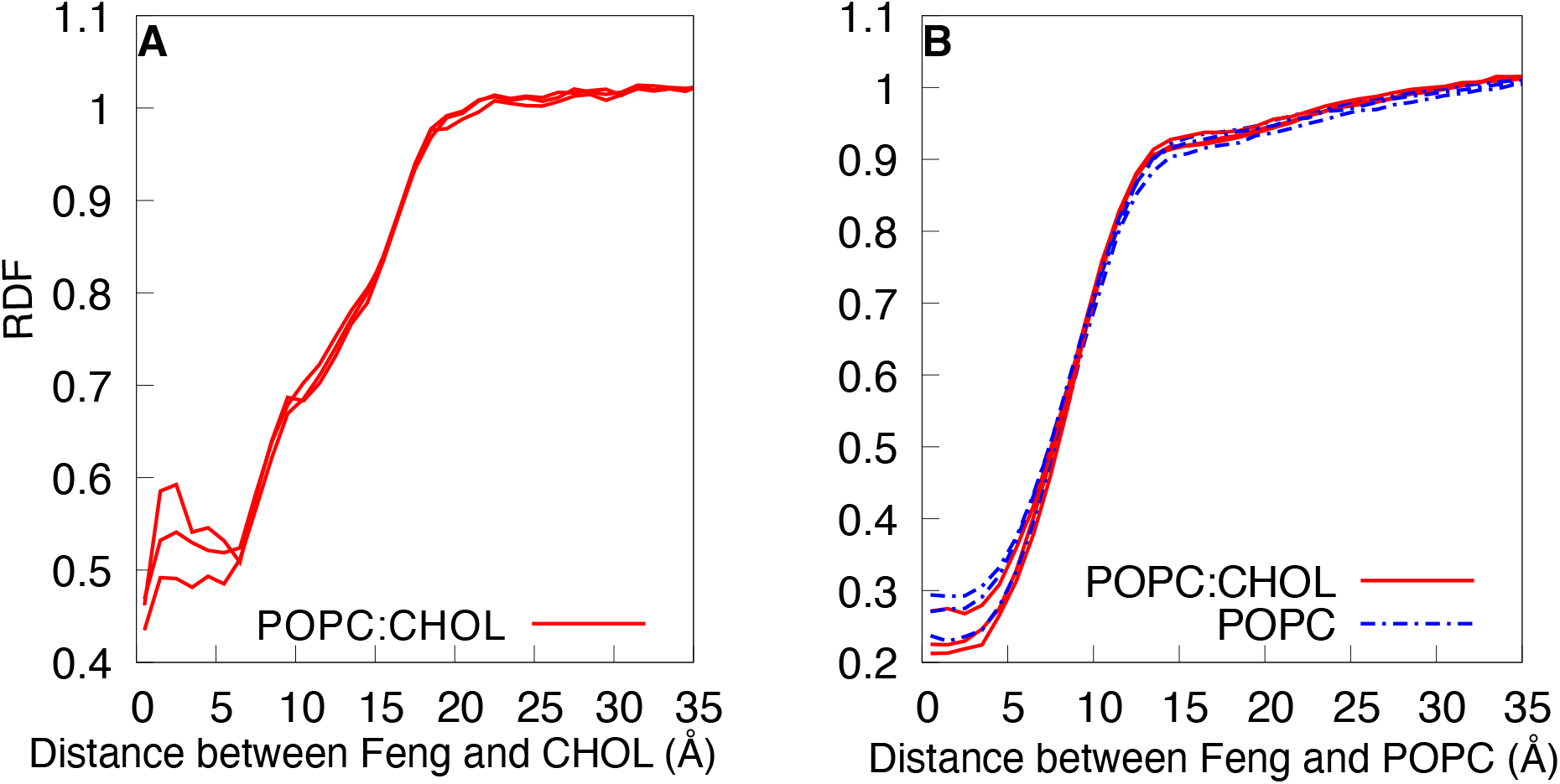
Lateral radial distribution function between A. cholesterol and fengycins. B. POPC and fengycins. Results from each weighted ensemble run are shown as individual curves.

Other than steric exclusion at short distances, there are no major features in either radial distribution function; if there were strong favorable interactions, one would expect to see enrichment at the contact distance, causing a peak from maybe 5-8 Å. This result is not consistent with the model proposed by Fiedler et al., who suggested that specific hydrogen bonds between cholesterol and fengycin could reduce fengycin aggregation.(31) These results are similar to our previous all-atom simulations, which also failed to show evidence for specific enrichment between POPC and fengycin.(3)

The only way fengycins could change the concentration of lipids at the surface would be by changing the area or creating more lipids. Another common hypothesis for the action of antimicrobial peptides is demixing of the membrane. To check for this, we computed radial distribution functions for cholesterol-cholesterol, POPC-POPC, and POPC-cholesterol in the presence and absence of fengycin; no enhanced sampling was needed to acquire good data for the pure membranes, so conventional molecular dynamics simulations were used (see Table 1).

Figure 6 shows that the cholesterol-cholesterol and POPC-POPC distributions are not significantly perturbed by fengycin; the curves with and without fengycin superpose almost exactly. However, there are some differences in the POPC-cholesterol distribution, within the first solvation shell. The presence of fengycin reduces the main contact peak — presumably because those contacts are replaced by fengycin — while there is a small increase in the population at very short distances. However, there is no perturbation beyond about 10 Å, and thus no effect on overall lateral structure of the membrane.

**Figure 6:**
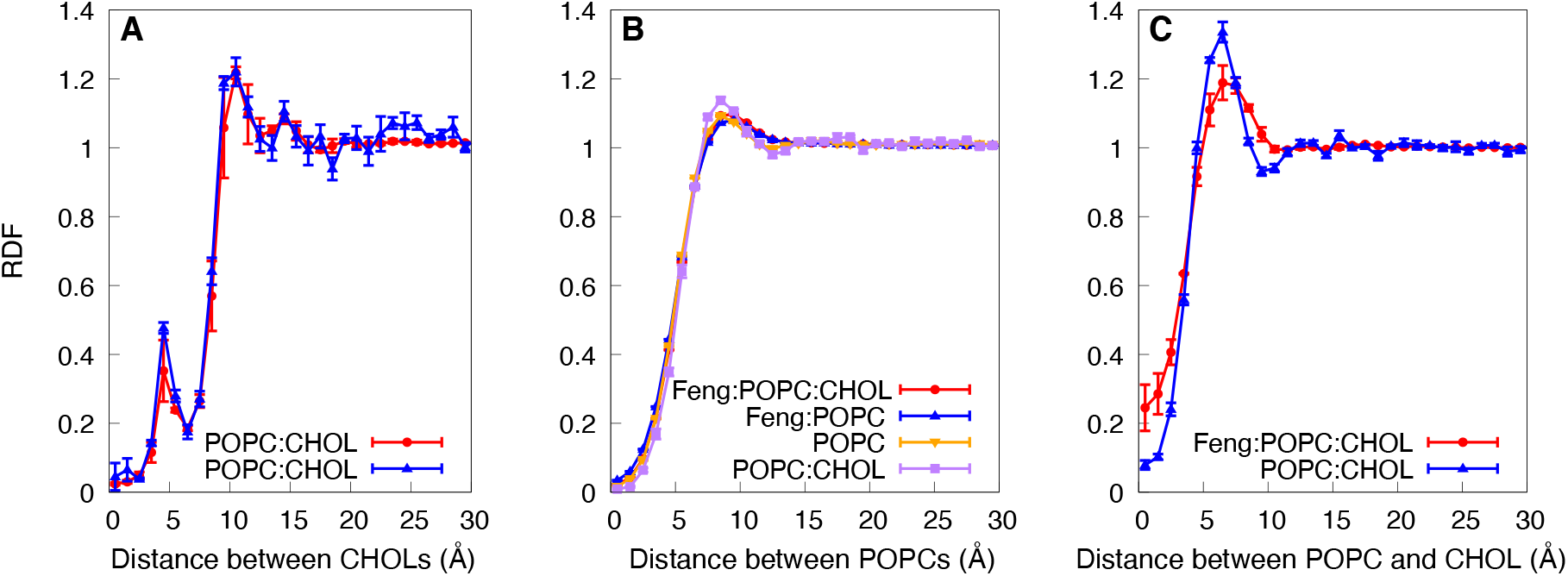
Lateral radial distribution function between A. cholesterols B. POPCs C. POPC and Cholesterol. All the lipid-lipid radial distribution functions are compared to their analogous value with no fengycins present.

### 3.4 Does cholesterol alter lipid packing around fengycin?

Since fengycin does not change the lateral organization of the membrane, we now consider an alternative hypothesis, that fengycin alters the local structure of the bilayer, specifically the lipid packing, in a cholesterol-dependent manner. To test this, we examine the 2D probability distribution for fengycin-fengycin and fengycin-lipid contacts, shown in Figure 7; Panels A and B show the results for the POPC and POPC:cholesterol bilayers, respectively, while Panel C shows the difference map.

**Figure 7:**
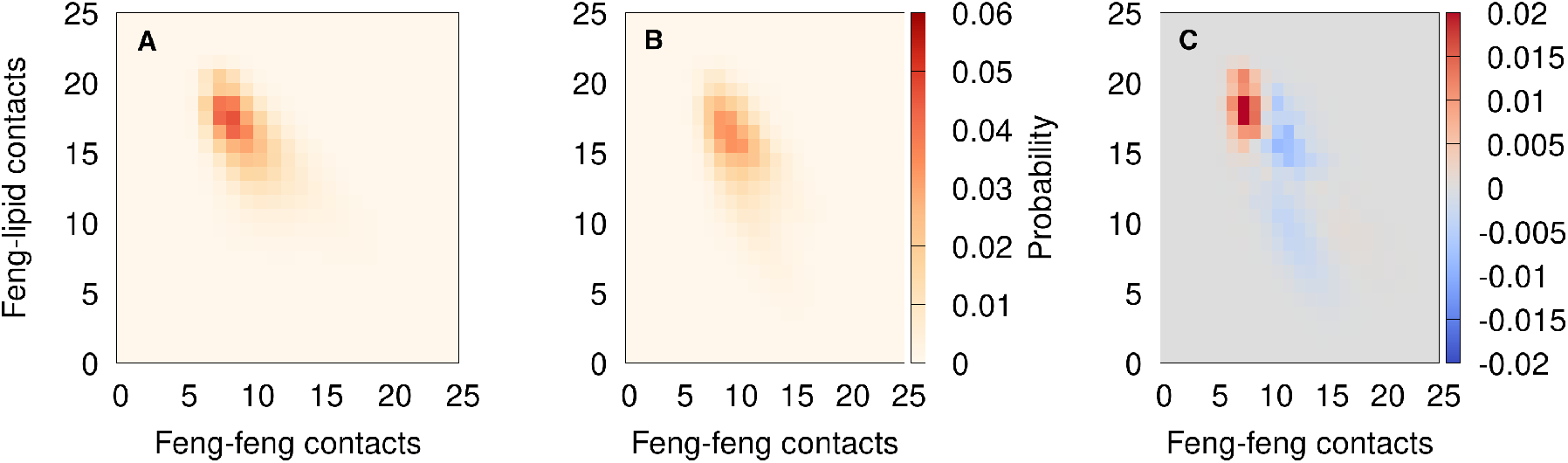
Probability of number of feng-feng contacts and number of feng-lipid contacts. A. POPC B. POPC-CHOL C. POPC-POPC:CHOL. We used equations 1 and 2 to calculate the different contacts.

While the two distributions are qualitatively similar, the POPC:CHOL distribution is broader, with the distribution shifted toward more fengycin-fengycin contacts and fewer fengycin-lipid contacts; this is most easily seen in Panel C.

### 3.5 Cholesterol limits fengycin’s disordering effects

Even without changing the lateral order of the membrane, fengycin can still induce leakage in a cholesterol-dependent manner by locally deforming or disordering the membrane. We quantify the chain packing using the molecular order parameter, a measure of whole-chain tilt (see Section 2.4.1 for details). Figure 8A shows the molecular order parameter for POPC saturated chains calculated as a function of lateral distance to the nearest fengycin. For comparison purposes, we also show the average result for the membranes with no fengycins added. The results show that the chain order is reduced at all distances in both systems, but unsurprisingly the effect is strongest close to the fengycin.

**Figure 8:**
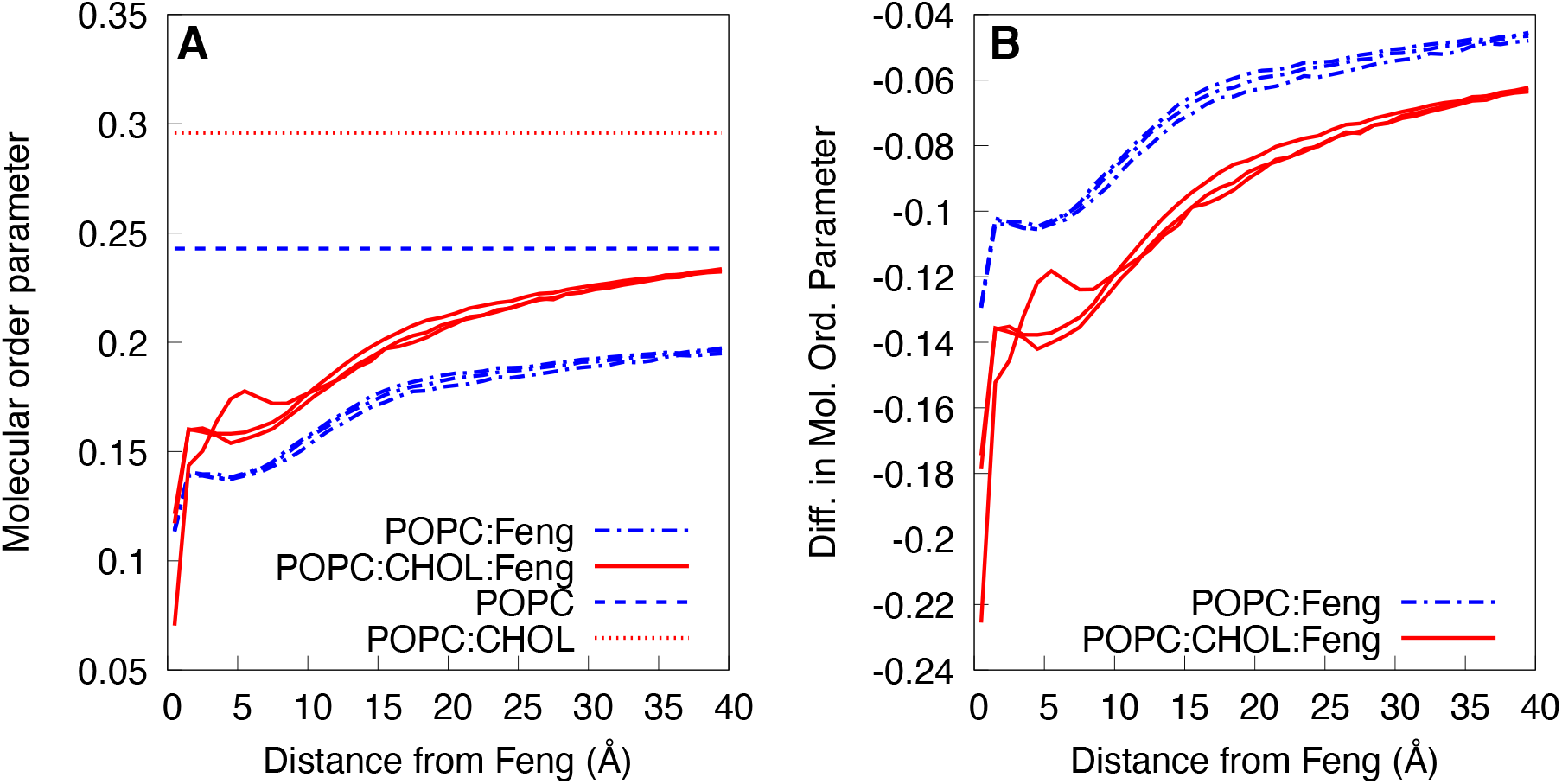
Molecular order parameters as a function of distance from nearest fengycin. A. Average molecular order parameters as a function of distance from fengycins. B.Difference between the distance-dependent molecular order parameter and the equivalent neat lipid system. Each curve shows the average in each for a single weighted ensemble replicate.

In Figure 8B, we have subtracted the bulk values (the dotted lines) from the distance-dependent curves, to simplify the process of quantifying difference in fengycin’s effect on the two systems, setting aside the fact that cholesterol already makes membranes thicker and more ordered. The curves computed for the POPC:cholesterol system are more negative at all distance ranges, which means that fengycin reduced the chain order of those membranes more than it did the neat POPC membranes, although overall the cholesterol-containing membranes are still more ordered.

To better understand this somewhat surprising result, we broaden this comparison to determine how the molecular order parameter simultaneously varies with distance and the aggregation state of the fengycin, shown as Figure 9. As expected, the same basic trend is observed: more fengycin-fengycin contacts implies more aggregation, which leads to lower molecular order parameters, particularly for those lipids close to the fengycin. However, when we examine the variation with distance, it’s clear that for any degree of aggregation, the reduced order parameters die off over shorter distances in cholesterol-containing membranes than in neat POPC.

**Figure 9:**
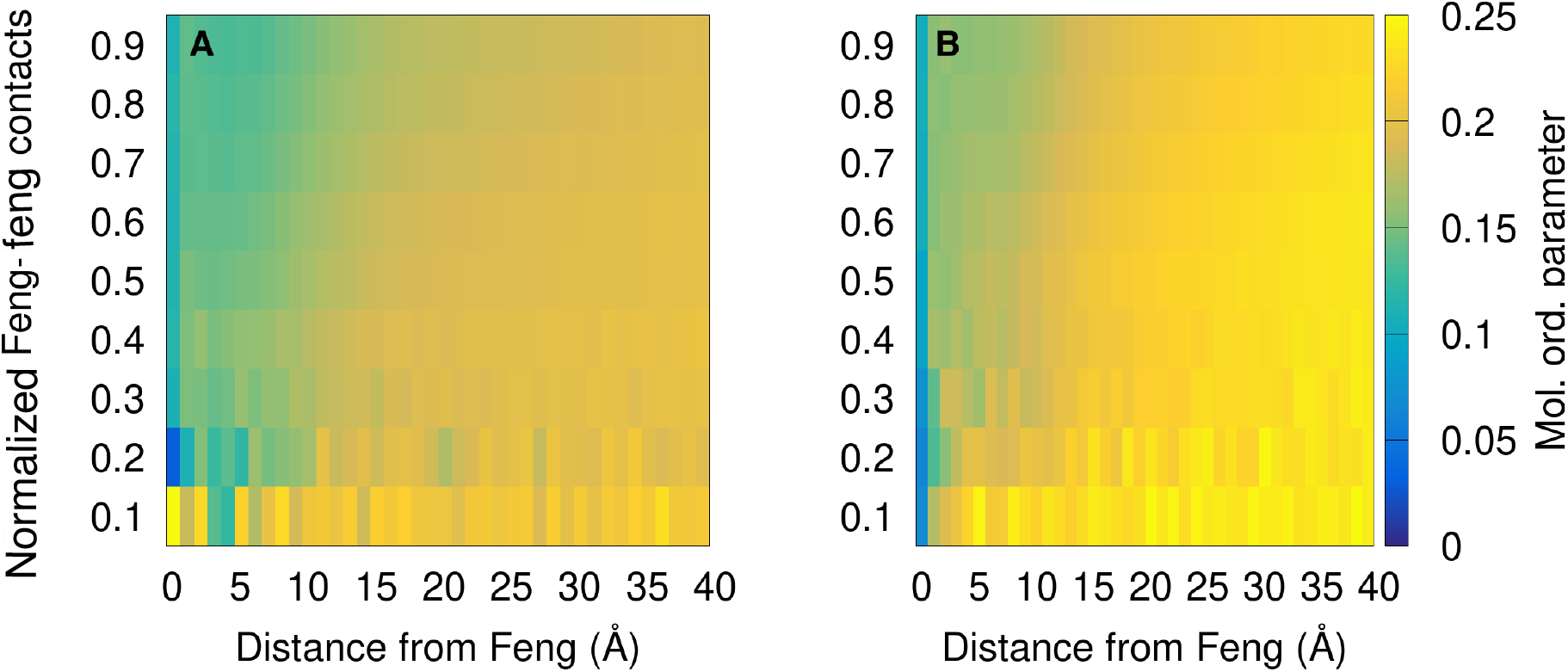
Average molecular order parameter for the palmitoyl chains as a function of distance from fengycin and fraction of fengycin-fengycin contacts. (A) POPC (B)POPC:CHOL. The x-axis is the lateral distance to the nearest fengycins and the y-axis is the fraction of the fengycin-fengycin contacts obtained by dividing the fengycin-fengycin contact with the highest measured fengycin-fengycin contacts. The color range indicates the molecular order parameter for the palmitoyl chains of POPC.

### 3.6 Fengycin bends the membrane

The previous coarse-grained simulations from Horn et al suggested that fengycin aggregates systematically bend the membrane when bound to one leaflet.(2) Visual inspection of the present simulations suggested similar behavior. To quantify this, we calculated the height of phospholipid heads — the distance to the center of the bilayer — with respect to the distance to the nearest fengycin, as discussed in Section 2.4.5.

To simplify comparison between membranes with difference thicknesses, we subtracted the average height of the phosphates from that leaflet in the neat simulations. The results are shown in Figure 10. Both compositions exhibit positive curvature; both leaflets are deflected in the +Z direction. Interestingly, the upper leaflets are distorted similarly, regardless of the composition of the surrounding membrane. By contrast, the lower leaflets’ response varies with cholesterol: the dynamic range for the lower leaflet’s deflection is 4 Å for POPC vs. 2.5 Å for the POPC:cholesterol mixture. The positive value for the lower leaflet suggests that the lower leaflet is pushed towards the membrane center. More the lower membrane is pushed towards membrane center, we are basically observing the thinning of the membrane. The overall amplitude of bending is most likely reduced by finite-size effects, since the bilayer contains only 186 lipids per leaflet.

**Figure 10:**
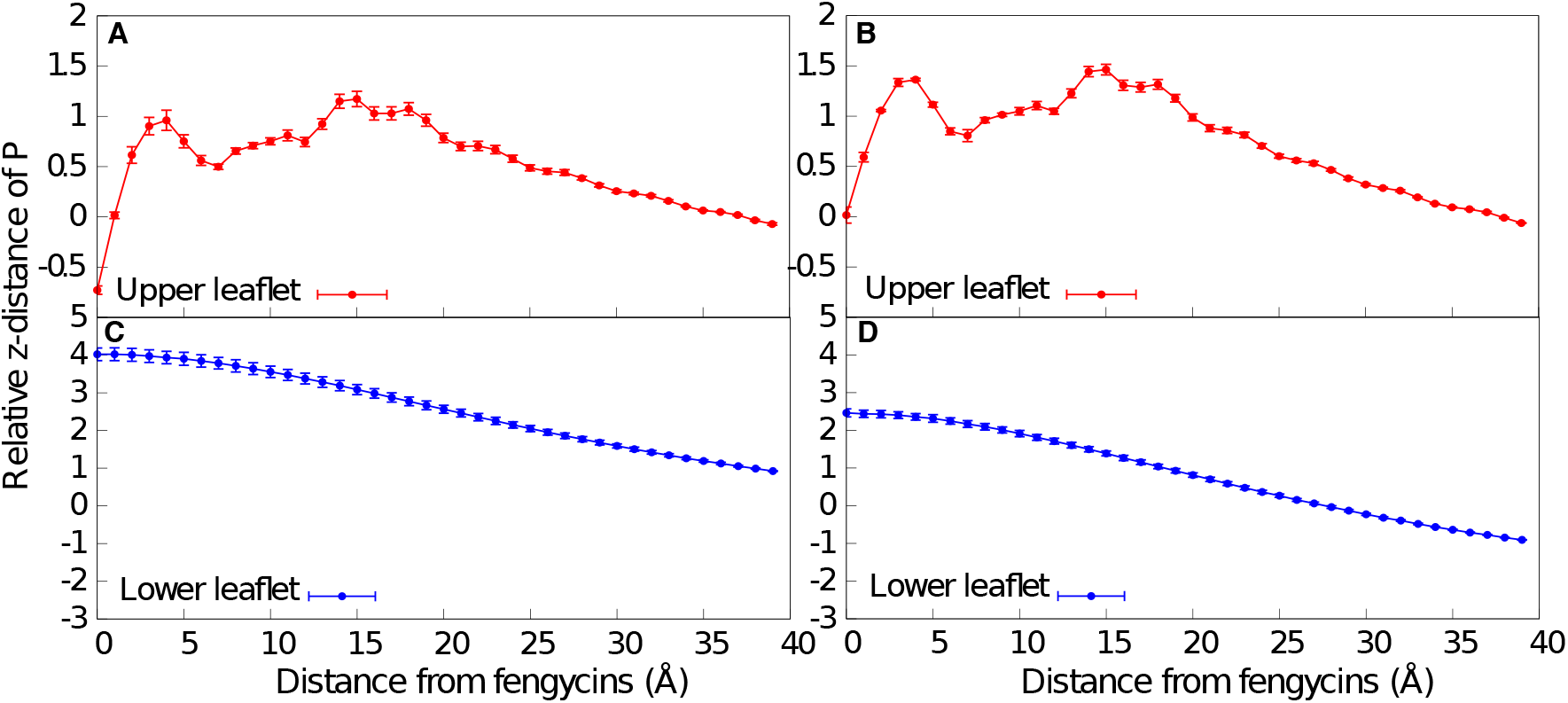
Relative Z-distances of phosphates for POPC (A and C) and POPC:CHOL (B and D) as a function of distance from fengycins. The x-axis is the distance of phospholipid from fengycin while y-axis denotes the height of phosphate moeity in the phospholipids after taking into account the thickness of the membrane. Each weighted ensemble run is considered as a single measurement and standard error is calculated using this assumption.

## 4 Discussion

The primary goal of the present work has been to quantitate the aggregation of fengycin in membranes and determine the role played by cholesterol in modulating that aggregation. This required developing a novel protocol to study the thermodynamics of peptide aggregation; in general previous simulation work focused only on its presence or absence (essentially, the sign of the Δ*G*). Here, we presented an enhanced sampling approach based on the weighted ensemble algorithm, which allowed us to detect subtle shifts in the aggregation behavior of fengycin in membranes with and without cholesterol.

As we were testing and developing a new method, we chose to perform the simulations using coarse-grained models, specifically the MARTINI coarse-grained force field, as opposed to fully atomistic models. This decision comes with certain trade-offs. One on hand, the coarse-grained models are computationally efficient, which was crucial to the success of the project. Lateral reorganization of the membrane is intrinsically a slow process, and even with enhanced sampling remains computationally expensive. At the same time, it was essential to run multiple independent calculations of each system, as this is the most reliable way to assess statistical uncertainties.

On the other hand, coarse-grained models by definition have less resolution than their atomistic counterparts, and that loss of resolution can at times affect both the research strategy and the results. For example, where mammalian membranes generally contain cholesterol, many fungal membranes often contain ergosterol. The present work focused on the affects of the presence or absence of cholesterol, but going forward with ergosterol is the next logical step, especially the recent work showing that ergosterol also modulates fengycin function. (49, 50) However, in the context of the MARTINI model cholesterol and ergosterol are very similar, so much so that distinguishing their effects appeared unlikely to succeed. We suspect that repeating these calculations with fully atomistic methods will be necessary.

The other major issue is the known tendency of the MARTINI ver 2.0 model to overstabilize protein-protein interactions. (32, 51, 52) We believe this manifests in the present work by causing the peptide aggregates to be more ordered than they were in our previous atomistic simulations;(3) forming stacked-disk aggregates maximizes protein-protein contact for cyclic peptides such as fengycin. Recently MARTINI ver 3.0 has been released which addresses this specific issue but due to the absence of cholesterol parameters for the same we were not able to run the simulation with the updated version.(53)

However, our earlier work showed that this effect is not overwhelming, in that some membrane compositions still did not lead to stable aggregates. (54) The main difference is that in those membranes there were favorable electrostatic lipid-peptide interactions that effectively competed with peptide-peptide packing, where in the present work we see no evidence for specific interactions between fengycin and either the POPC headgroups or cholesterol.

Lipopeptides such as fengycin are unlikely to exist as monomers in solution; rather, one would expect to see the formation of micelles that can bury the acyl chain while leaving the relatively polar peptide portion exposed to water. Previous work from our group on a difference family of lipopeptides (55) showed that differences in the kinetics of binding from a micellar state provided an alternative explanation for membrane selectivity. These were exceptionally challenging calculations, but it would be interesting going forward to determine if analogous phenomena are relevant here, and in particular whether cholesterol and/or ergosterol modify the binding kinetics of fengycin micelles.

## 5 Conclusions

Fengycins are antifungal lipopeptides that permeabilize fungal membranes while leaving the bacterial species synthesizing it unharmed and causing minimal damage to mammalian cells. The present work is a step toward understanding their mechanism, which characterizes the propensity to aggregate and how that aggregation is modulated by cholesterol. We conclude that cholesterol subtly shifts the aggregation propensity, and more significantly that the presence of cholesterol reduces the range over which fengycin perturbs the membrane. These actions are not the result of direct specific interactions between fengycin and cholesterol, but rather reflect the balance of interactions between all membrane constituents.

## Acknowledgements

We would like to thank the Department of Biochemistry and Biophysics at the University of Rochester Medical Center for their support throughout this project. We also appreciate the computing resources provided by Center for Integrated Research Computing at the University of Rochester.

